# Differential gene expression underpinning production of distinct sperm morphs in the wax moth *Galleria mellonella*

**DOI:** 10.1101/2023.12.13.571524

**Authors:** Emma Moth, Fiona Messer, Saurabh Chaudhary, Helen White-Cooper

**Author notes:** Current address: Wellcome Trust-Medical Research Council Cambridge Stem Cell Institute, University of Cambridge, Cambridge, UK.

## Abstract

Male Lepidoptera makes two distinct sperm types; each ejaculate contains both eupyrene sperm, which can fertilise the egg, and apyrene sperm, which are not fertilisation competent. These sperm have distinct morphologies, unique functions, and different proteomes. Their production is highly regulated, however very few genes with specific roles in production of one or other morph have been described. We present the first comparative transcriptomics study of precursors of eupyrene and apyrene sperm to identify genes potentially implicated in regulating or enacting the distinct differentiation programmes. Differentially expressed genes included genes with potential roles in transcriptional regulation, cell cycle and sperm morphology. We identified gene duplications generating paralogues with functions restricted to one or other morph. However phylogenetic analysis also revealed evolutionary flexibility in expression patterns of duplicated genes between different Lepidopteran species. Improved understanding of Lepidopteran reproduction will be vital in targeting prevalent pests in agriculture, and on the flip side, ensuring the fertility and thus survival of pollinator populations in response to environmental stress.

## Introduction

Spermatogenesis is a highly regulated process that results in the generation of mature sperm with highly specialised morphology. In insects, the process typically involves continued production of sperm throughout the adult life of the male, sustained by a male germline stem cell population. Stem cell divisions generate spermatogonia committed to differentiation, which then undergo mitotic amplification before differentiating into primary spermatocytes and switching to a meiotic cell cycle. Post-meiotic morphological changes include spermatid elongation, nuclear reshaping, mitochondrial reorganisation, and growth of the axoneme. Each spermatogonium is enveloped by somatic cyst cells that form a squamous epithelium within which the germline cells differentiate. In Lepidoptera each male makes two distinct sperm types, a phenomenon known as sperm heteromorphism (see (1, 2) for detailed reviews). Each ejaculate contains both fertilising eupyrene sperm and non-fertilising apyrene sperm. Eupyrene spermatogenesis initiates first, during larval life. In early pupal development a hormonally driven switch intiates apyrene spermatogenesis (3).

Eupyrene and apyrene sperm are morphologically very distinct. Eupyrene sperm are longer, they remain bundled together in the ejaculate, only becoming separate within the female genital tract (4). The shorter, apyrene, sperm lack both a nucleus and an acrosome, explaining their inability to fertilise eggs. Both sperm morphs are motile, and this motility is required for normal reproduction (5, 6). Consistent with these dramatic differences in the final sperm, the process of spermatogenesis differs between the two morphs. Early primary spermatocytes are bi-potential; those in larval testes proceed along the eupyrene differentiation programme. Early pupae produce an as yet unidentified hormone, termed ‘apyrene spermatogenesis inducing factor’ (ASIF). On receipt of ASIF, spermatocytes that are still bi-potential switch programme and eventually differentiate into apyrene sperm. Already committed eupyrene spermatocytes do not respond to ASIF, and remain on their eupyrene developmental trajectory. (1). Despite their different potential, primary spermatocytes on the two pathways are not morphologically distinguishable until they initiate the meiotic divisions.

Production of eupyrene spermatocytes involves a conventional meiosis I spindle, with robust microtubule arrays, a well-formed metaphase plate, and segregation of homologous chromosome in anaphase I (7). In contrast, the meiosis I spindle in cells destined to become apyrene sperm is much less robust, with reduced microtubule arrays and a poorly defined metaphase plate. There is extensive chromosome non-disjunction leading to aneuploid cells with dispersed micronuclei. As eupyrene spermatids elongate, the nuclei cluster at one end of the cyst, become needle-shaped and intimately associate with the overlying cyst cell. In contrast, micronuclei in apyrene spermatids cluster in the middle of the elongating cyst and are gradually degraded (8). Spermatid individualisation of both morphs involves peristaltic squeezing of cyst cells; removing excess cytoplasm from all spermatids, and forcing elimination of nuclei from apyrene spermatids.

The final product of this deliberate and orchestrated process is two morphs with different morphologies made with different proteomes, comprising some shared proteins and some proteins unique to one or other morph (9, 10). A small number of genes have been demonstrated via CRISPR-Cas9/RNAi experiments to be important for Lepidopteran sperm heteromorphism, including *Sex-lethal (Sxl)* (Table S1) (5, 6, 11, 12). However, there has not been a systematic, unbiased identification of differentially expressed genes that ensure normal differentiation of spermatocytes towards eupyrene and apyrene sperm fates.

To identify genes that may be required for the alternative differentiation trajectories we compared the transcriptomes of spermatocytes from larval *Galleria mellonella* testes, that were destined to become eupyrene sperm, with spermatocytes from pupal testes that were destined to become apyrene sperm. A high number of differentially expressed genes were found, including transcription factors, meiotic regulators and sperm axoneme components.

Comparison of our transcriptomic dataset with mature sperm proteomic data from other lepidoptera (10), and subsequent phylogenetic analysis, enabled validation of our RNA-seq data, as consistent with the resulting mature sperm proteomes. Furthermore, phylogenetic analysis elucidated the evolution of eupyrene– and apyrene-enriched paralogues of both the sperm axoneme component *Ccdc63*, and β*-tubulin* in moths, providing an insight into the evolution of Lepidopteran sperm heteromorphism.

## Materials and Methods

### G. mellonella culture

Wild type *G. mellonella* larvae were provided by the Galleria mellonella Research Centre (Exeter University) and initially stored in the dark at room temperature and maintained on food medium based on diet 3 of (13) (Supplementary methods). Larvae were subsequently incubated at 30°C to induce pupation (13).

### Testis spills

Individual follicles were dissected from three last instar larval testes and three pupal testes, cut open in 10 µl PBT (1x PBS, 0.1% Tween 20) and pipetted onto a slide. Paraformaldehyde (10µl of 4% w/v in PBT) was added for 10 minutes at room temperature, and then 1 µg/ml Hoescht (33258) in mounting medium (2.5% n-propyl gallate in 85% glycerol,) was added. Fluorescence was analysed using Olympus Bx50 and images taken with a Hamamatsu ORCA-05G camera and HCImage software.

### Fluorescence Hybridisation Chain Reaction (HCR-FISH)

HCR-FISH v3 (14) was used to visualise mRNA expression in *G. mellonella* larval (n=10) and pupal (n=10) testes (14) (See Supplementary Methods 2 for detailed protocol) using 2-4 oligonucleotide probe pairs per gene (Supplementary Table 2). Fluorescence was visualised using either the Zeiss Lightsheet Z.1 system or the Zeiss LSM880 Airyscan upright confocal microscope in the Cardiff Bioimaging Hub (Supplementary methods).

### RNA-seq of *G. mellonella* primary spermatocyte cysts

Two primary spermatocyte cysts were collected from each of five 6^th^ instar larvae and five three-day old pupae (Supplementary Methods). RNA libraries were produced from the 20 primary spermatocyte cysts using the QIAseq FX Single Cell RNA Library Kit (Qiagen). Library quality and fragment size was assessed with the D1000 Tapestation (Agilent) and DNA size selection with the Blue Pippin system (Sage Science), by the Cardiff Genomics Research Hub. Libraries were sequenced on an Illumina NextSeq500 Sequencer (Supplementary methods).

### Bioinformatics and statistical analysis

A standard RNA-seq bioinformatics pipeline was used to assess sequence quality, align reads to the *G. mellonella* reference genome (CSIRO_AGI_GalMel_v1, Rahul Vivek Rane 2022) from NCBI (15) and count reads that mapped to genomic features (Supplementary methods 5, Table S3). Statistical analysis was carried out in R studio. SARTools R package (16) with DEseq2 (v1.38.3) was used for normalisation of the data and differential gene analysis. Samples with low reads and/or ambiguous larval vs pupal clustering after Principal Component Analysis (PCA) and hierarchal clustering, were removed before final DEseq2 analysis. Heatmaps were created using normalised counts of top 100 upregulated DEGs and top 100 downregulated DEGs using the ComplexHeatmap (v2.14.0) package with Pearson clustering (17). All significant (p<0.05) DEGs were input into DAVID Bioinformatics database gene conversion tool to obtain gene names (18). DAVID Functional annotation analysis was completed to obtain enriched GO and KW terms (p<0.2).

### Phylogenetic analysis of DEGs of interest

G. mellonella protein sequence of interest was input into BLASTP as a query sequence against Galleria mellonella, Bombyx mori, Manduca sexta, Danaus plexippus, Drosophila melanogaster, Aedes aegypti, Homo sapiens proteomes. Phylogenetic analysis was then completed using MEGA V. 11.0. 13 software (19) (Supplementary Methods). Resulting phylogenetic trees were then cross-referenced with our RNA-seq dataset and previously published proteomic and transcriptomic datasets (10, 20–23).

## Results

### Validation of the switch to apyrene sperm development in *G. mellonella* pupae

To validate the switch to apyrene sperm production after the onset of pupation, individual testis follicles from *Galleria mellonella* 6th instar larvae and 3-day old pupae were stained for DNA. Whole testes and spilled testis contents confirmed eupyrene spermatogenesis in larval stages (Fig. 1A, C, E) and apyrene spermatogenesis in pupal stages (Fig. 1B, D, F). Primary spermatocytes were morphologically indistinguishable between larval and pupal testes (Fig. 1C, D, large arrows). Early haploid eupyrene spermatids were observed in larval testis (Fig. 1C, small arrow), with spermatids subsequently completing spermiogenesis in pupal stages to form bundles with elongated nuclei (Fig. 1D, E small arrowheads). Importantly, early apyrene spermatids with centrally located nuclei were only observed in pupal testes (Fig. 1D, E, large arrowheads). Therefore, we concluded that primary spermatocytes collected from larvae and pupae were representative of eupyrene and apyrene development, respectively.

**Figure 1.**
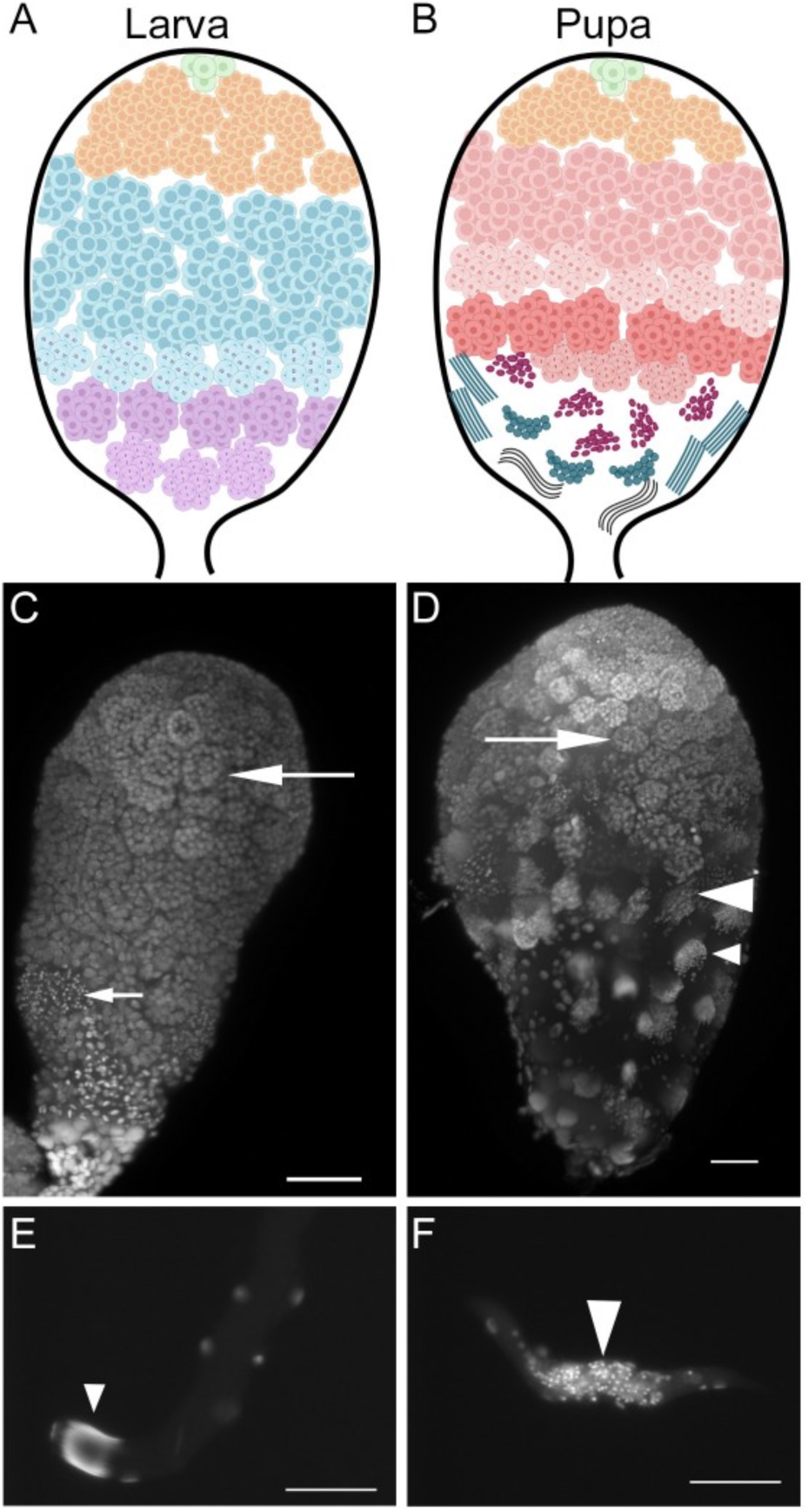
*G. mellonella* larval and pupal testis morphology. Schematic representations, and whole mount testes stained for DNA, of larval (A, C) and pupal (B, D) testes from *G. mellonella*. Germline stem cells and spermatogonia reside at the apical tip (green), and generate cysts of bipotential early spermatocytes (orange) encapsulated by cyst cells. In larvae these differentiate into late primary spermatocytes on the eupyrene sperm trajectory (blue in A, large arrow in C), which undergo meiosis and become secondary spermatocytes and then early spermatids (purple in A, small arrow in C). In pupal testes late spermatocytes (pink in B, large arrow in D) differentiate along the apyrene sperm trajectory, to generate secondary spermatocytes and spermatids (dark pink in B, large arrowhead in D). Pupal testes also contain a few eupyrene secondary spermatocytes (small arrow in D) and spermatids (teal in B, small arrowhead in D). DNA staining of eupyrene spermatid (E) and apyrene spermatid (F) cysts, with nuclei at the end of the cyst (small arrowhead) or centrally located (large arrowhead) respectively. Scale bar is 50μm.

### *Gmsxl* expression persists longer in cells progressing though the apyrene differentiation pathway

The RNA-binding protein Sxl is required for apyrene sperm development in *Bombyx mori,* but for development of both sperm morphs in the tobacco cutworm, *Spodoptera litura* (5, 6, 24). In *B. mori*, *Bmaly*, a homologue of the *D. melanogaster* meiotic transcriptional regulator *aly* and its paralogue *lin9*, is required for progression of spermatocytes into the meiotic divisions in larval testes; its role in pupal testes has not been evaluated (25). To evaluate expression of these genes in both eupyrene and apyrene differentiation in *G. mellonella* we used HCR-FISH on larval and pupal testes.

*Gmlin9* was expressed through both developmental trajectories. An abrupt increase in expression was found as spermatocytes matured in larval testes (Fig. 2A). In pupal testes *Gmlin9* expression increased steadily, peaking in mature primary spermatocytes (Fig. 2B). *Gmlin9* transcript gradually declined through secondary spermatocytes and spermatids (Fig. 2 A, B, small arrowheads). In larval testes, *Gmsxl* expression was high in late spermatogonia and early primary spermatocytes (Fig. 2C, G, small arrows), and the transcript abruptly declined in late spermatocytes (Fig. 2C, G, large arrows). No signal was detected in eupyrene cysts undergoing meiotic divisions (Fig. 2C, small arrowhead)). Thus, cysts on the eupyrene developmental trajectory initially had high *Gmsxl* and low *Gmlin9* before switching to a low *Gmsxl*, high *Gmlin9*, state as late spermatocytes.

**Figure 2.**
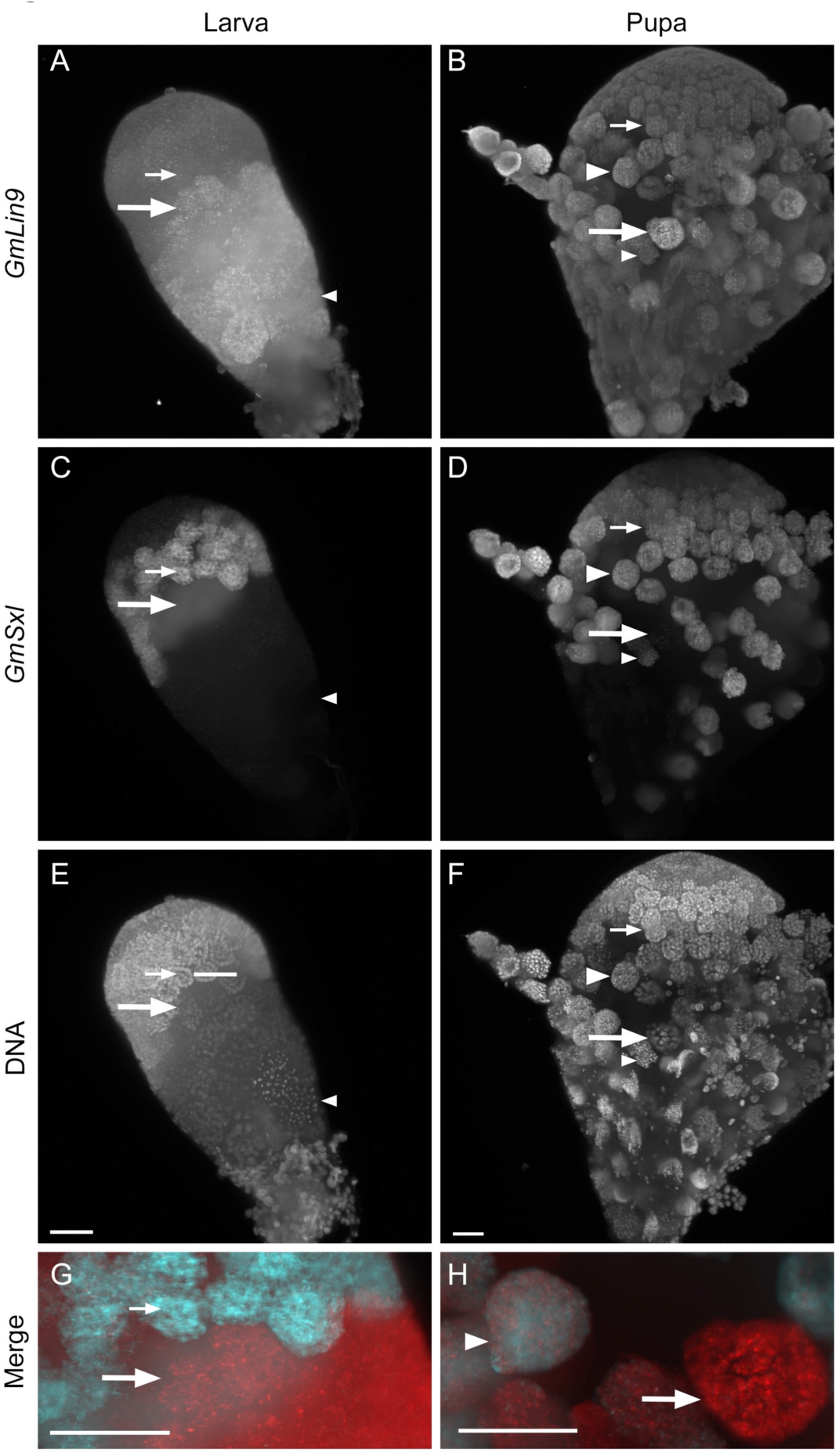
*Gmsxl* expression persists longer in the apyrene spermatogenesis programme. HCR-FISH analysis of Gmlin9 (A, B, red in merged images, G, H) and *Gmsxl* (C, D, cyan in merged images G, H), and DNA staining (E, F) of larval (A, C, E, G) and pupal (B, D, F, H) testes. *Gmlin9* expression is low and *Gmsxl* expression is high in early spermatocytes (small arrows). Spermatocyte cysts that will differentiate into eupyrene sperm have high *Gmlin9* and low *Gmsxl* (large arrow). Spermatocyte cysts that will differentiate into apyrene sperm have high *Gmlin9* and high *Gmsxl* (large arrowhead). Imaged using the Zeiss Lightsheet Z.1 system. Scale bar is 50μm.

In pupal testes *Gmsxl* was expressed in late spermatogonia and early spermatocytes (Fig. 2D, small arrow) and was detected at high levels in many late primary spermatocyte cysts (Fig. 2D, H, large arrowhead). A few very late spermatocytes lacked *Gmsxl* transcript (Fig. 2D, H, large arrow); these are likely to be cysts that had already committed to eupyrene differentiation before the early pupal action of ASIF. Secondary spermatocytes and spermatids destined to become apyrene sperm also retained some *Gmsxl* transcript (Fig. 2D, small arrowhead). Thus, cysts on the apyrene developmental trajectory initially had high *Gmsxl* and low *Gmlin9*, then had high *Gmsxl* and high *Gmlin9* before switching to a low *Gmsxl*, medium *Gmlin9*, state as early spermatids.

### RNA-seq of primary spermatocytes from larval vs pupal testes

The differential expression of *Gmsxl* in spermatocytes on the two differentiation pathways confirms that these morphologically identical cells have different transcriptome profiles. We used an unbiased approach to investigate transcriptomic differences between primary spermatocyte cysts destined to become eupyrene sperm vs apyrene sperm. RNA-seq was conducted on individual primary spermatocyte cysts collected from larval (10 cysts) and pupal testes (9 cysts) (Fig S1). Results for trimming and subsequent mapping to the *G. mellonella* reference genome are shown in Supplementary Table 3. Larval sample 5.2 was excluded at this stage due to low mapping percentage (Table S3).

### Larval and pupal cysts clustered separately in Principal Component Analysis plots

The morphological examination and *Gmsxl* FISH both indicated that day 3 pupal testes contain a few late spermatocyte cysts that are on the eupyrene sperm differentiation pathway, having been early primary spermatocytes just past the commitment point when the early pupal ASIF induced switch occurred. To ensure unambiguous eupyrene and apyrene samples for valid differential gene expression (DEG) analysis, hierarchal clustering and principal component analysis (PCA) were conducted. Six larval samples and seven pupal samples were included in the final transcriptomic analysis. Two larval samples (2.1 and 5.1) clearly clustered away from other larval samples (Fig. 3A), whilst another larval sample (1.1) clustered within pupal samples in hierarchal clustering (Fig. S2A). Pupal sample 2.1 clustered very closely to larval samples (Fig. 3A), and was potentially a eupyrene-destined primary spermatocyte. Pupal sample 2.2 also clustered away from other pupal samples when comparing PC1 and PC3 axes (Fig. S2A). Therefore, these samples were removed from final DEG analysis to ensure a biologically valid comparison (Fig. S2B). DEG analysis using DESeq2, with an adjusted p-value threshold of <0.05, identified 373 genes significantly upregulated in primary spermatocytes from larvae, and 686 genes significantly upregulated in pupal primary spermatocytes (Fig. 3B, Supplementary Data Files 1, 2 & 3). While we did not impose a fold change cut off, in practice all bar one of these differentially expressed genes showed a 2-fold or more difference between cyst types.

**Figure 3.**
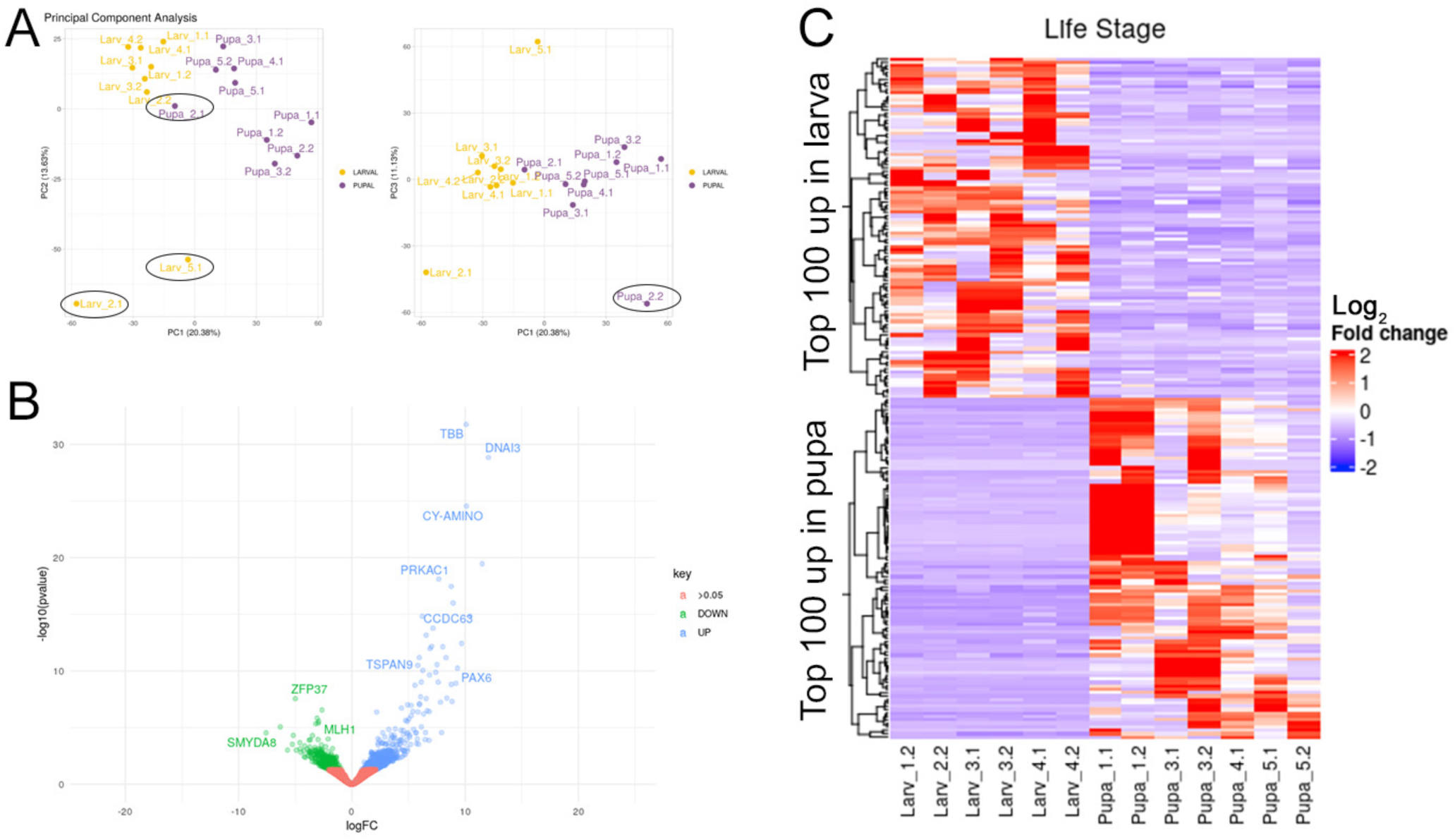
RNA-seq of larval and pupal spermatocytes reveals differential transcriptomes. A) Principal component analysis of cyst sequencing. Cysts excluded from further analysis are circled. B) Volcano plot of all detected genes comparing expression in larval cysts with pupal cysts reveals more genes significantly upregulated in pupal cysts. C) Heat map of genes with highest fold changes, revealing the variability in signals in the different cysts comprising the whole sample.

### DAVID Functional Enrichment analysis highlighted genes involved in core biological processes in spermatogenesis

The DEG lists were input into DAVID Functional Enrichment analysis (see Supplementary Data File 4 for full lists). Terms such as ‘Transcription’, ‘DNA binding’, ‘zinc ion binding’ were enriched in larval spermatocytes, whilst ‘Cell division’, ‘Motile cilium’, ‘Coiled-coil’ were enriched in pupal spermatocytes (Supplementary Data File 4). These terms are biological processes and protein properties expected to appear in sperm development, and there is no obvious enriched term to explain the fertile vs infertile sperm fate decision in early spermatocytes. The genes enriched in the gene ontology (GO) and keywords (KW) terms were then investigated further via literature analysis. Several were previously known to be involved in spermatogenesis, summarised in Table S4. Overall, our DEG analysis has revealed many genes of interest involved in core biological processes in spermatogenesis that are differentially expressed between the early eupyrene– and apyrene-committed cells, with a large scope for future exploration.

We examined the RNA-seq data for *Gmlin9* and *Gmsxl* genes, to evaluate if the HCR-FISH and RNA-seq data were consistent. As expected from HCR-FISH analysis, *Gmlin9* (LOC113523285) was not differentially expressed between larval and pupal spermatocytes (Fig. 2, Table S4, S5). Interestingly, *Gmsxl* (LOC113515001) was upregulated in pupal spermatocytes, supporting the HCR-FISH finding that *Gmsxl* expression persists to a later stage in apyrene sperm development (Fig. 2, Table S5). However, this upregulation was not significant (Log2FC= 1.475, P.adj = 0.081399). Overall, the corroboration of *Gmsxl* and *Gmlin9* HCR-FISH expression patterns in *G. mellonella* testes and RNA-seq expression values validates the predictive value of our RNA-seq dataset.

We also investigated RNA-seq results for other previously discovered sperm heteromorphism regulators in Lepidoptera (Table S1). None were differentially expressed between larval and pupal spermatocytes. Interestingly, two genes implicated in sperm heteromorphism regulation in *B. mori* (*Maelstrom* and *PNLDC1*) were barely detected in sequenced *G. mellonella* primary spermatocyte cysts, suggesting differences between Lepidopteran species (Table S1).

### *GmTaf4* is expressed at higher levels in larval than pupal primary spermatocytes

Among the DEGs contributing to the ‘Transcription’ annotation term enrichment in larval spermatocytes was LOC113519479, which encodes Taf4, a subunit of the general transcription factor complex TFIID. In *D. melanogaster* a testis-specific paralogue of *Taf4*, *nht*, is critical for testis-specific transcription (26). BLAST searches confirmed that this is a single copy gene in *G. mellonella* and other sequenced Lepidoptera. A difference in expression of this gene could result in higher total transcriptional activity in larval spermatocytes compared to pupal spermatocytes. HCR-FISH showed that *taf4* is expressed in primary spermatocytes in both larval and pupal testes, as expected given its critical role in transcription, but also confirmed that the transcript was more abundant in late larval spermatocytes than pupal spermatocytes at the same differentiation stage (Fig. 4).

**Figure 4.**
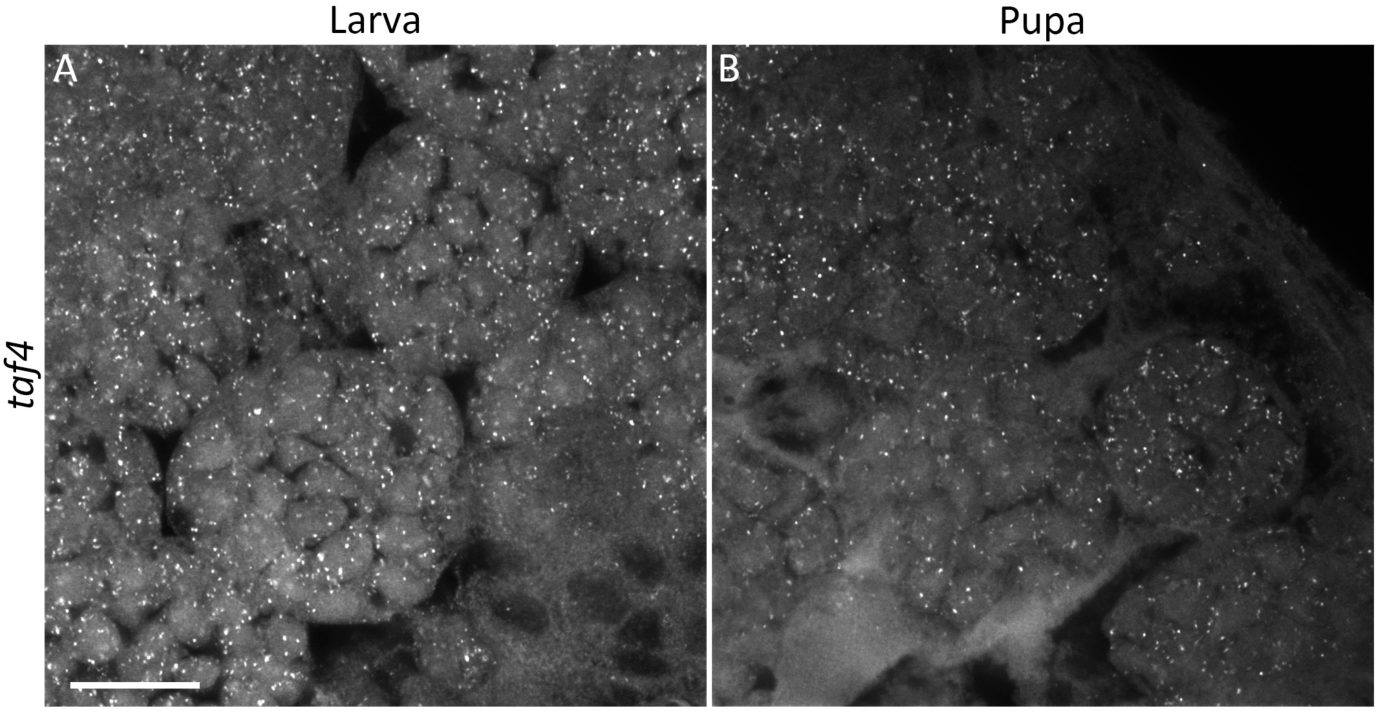
*Taf4* expression is higher in larval spermatocytes than pupal spermatocytes. HCR-FISH analysis of *GmTaf4* in larval (A) and pupal (B) spermatocytes. Imaged using the same acquisition settings for both samples on the Zeiss LSM880 Airyscan upright confocal microscope. Scale bar is 20μm.

### Gene duplication and specialisation has produced apyrene– and eupyrene-enriched *Ccdc63* paralogues in moths

One of the most dramatically upregulated genes in pupal spermatocytes was coiled-coil domain containing protein 63 (*Ccdc63,* LOC113513840) (Fig. 3B, Table S4). Ccdc63 is a component of the outer dynein arm docking complex, involved in formation of the sperm axoneme. Phylogenetic analysis of *Ccdc63* evolution, incorporating Lepidopteran and Dipteran species, revealed a series of gene duplication and sub-functionalisation events, to produce somatic-, germline-, and morph-enriched paralogues. In Diptera, duplication of the ancestral gene generated a somatically expressed paralogue and a germline expressed paralogue. In *D. melanogaster*, these are *Ccdc114* (*CG14905*), expressed in Johnston’s organ neurons which possess motile cilia, and *wampa* (23, 27), which encodes a component of the sperm proteome (28) respectively. In *A. aegypti,* AAEL011965 (LOC5575638) is highly expressed in the antenna (29) while the *wampa* orthologue, AAEL007188 (LOC5568877) is highly expressed in the testis (20, 21). In Lepidoptera, a similar but independent duplication of the ancestral gene generated somatic– and germline-enriched paralogues (Fig. 5). Interestingly, the post-duplication germline gene underwent a further duplication to give two germline enriched paralogues in all Lepidopteran species analysed. *M. sexta* sperm proteomic data revealed specialisation of one germline paralogue for eupyrene sperm, and the other paralogue for apyrene sperm (10). The *M. sexta* apyrene-enriched protein (LOC115451629) clustered in the phylogenetic tree with *G. mellonella Ccdc63* (LOC113513840), which was highly expressed in apyrene-destined spermatocytes (red cluster, Fig. 5). Further evidence to support the specialisation of these paralogues in moths is provided by transcriptomic data from *B. mori* larvae which detected enrichment of predicted eupyrene-enriched *Ccdc63* (LOC101738429) transcripts in larval testes, but not *Ccdc63* (LOC101746125) transcripts predicted to be apyrene-enriched (higher expression in pupal testes) (22). This apyrene specialisation may be exclusive to moths, as the paralogous protein in the Monarch butterfly *D. plexippus* (LOC116774756) was found to be enriched in eupyrene sperm, rather than apyrene (10). Based on the name of the *D. melanogaster* homologue, *wampa* (27), we named the largely apyrene-enriched paralogue (LOC113513840) *wimpa*, and the eupyrene-enriched paralogue (LOC113515144) *wompa*.

**Figure 5.**
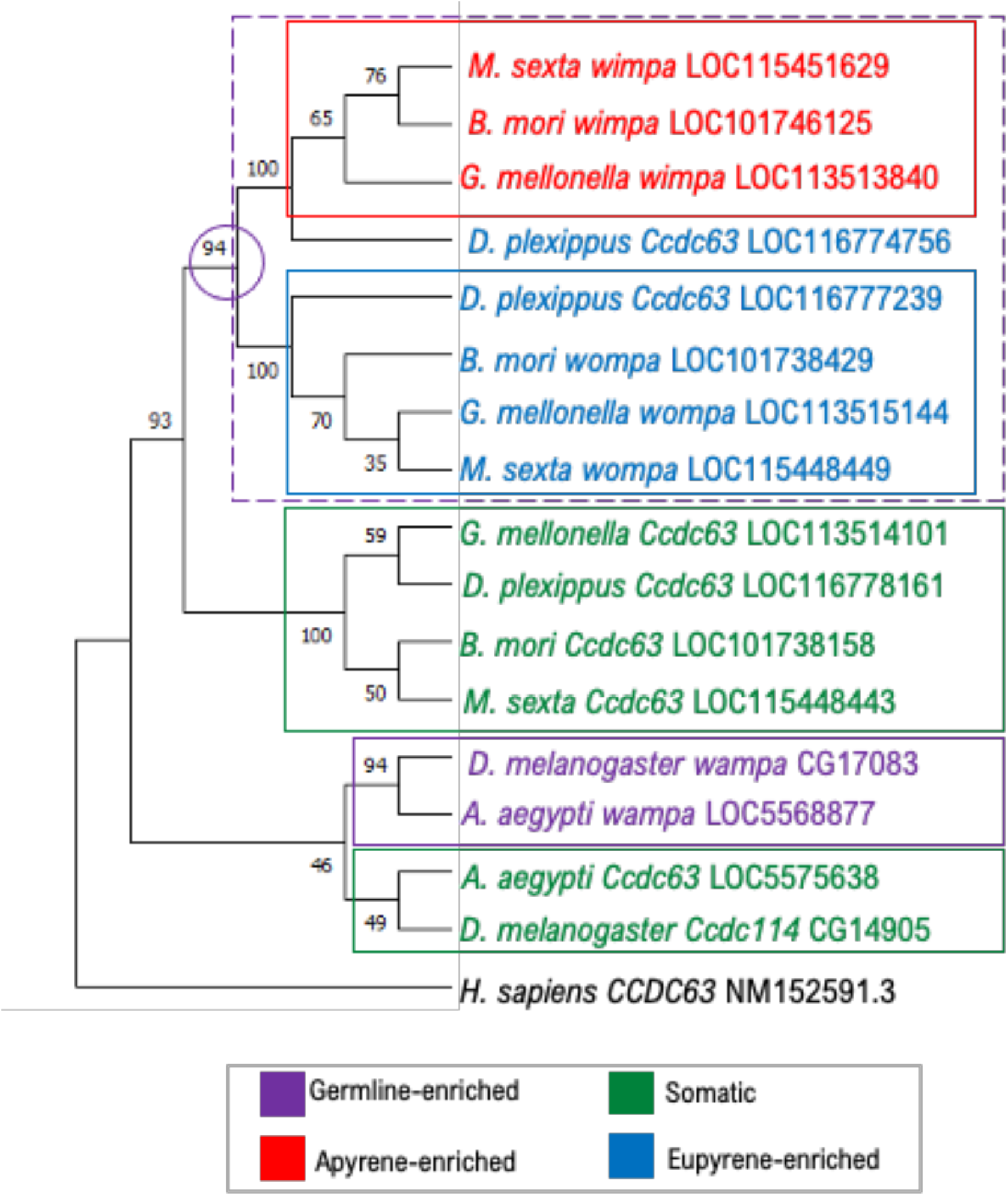
*Ccdc63* phylogenetic tree reveals gene duplication and subfunctionalisation events in Lepidoptera. Maximum likelihood tree of conserved Ccdc63 protein regions. 16 Ccdc63 homologues were analysed from Lepidopteran (*G. mellonella, D. plexippus, M. sexta, B. mori*) & Dipteran (*D. melanogaster, A. aegypti*) species, with human (*H. sapiens*) CCDC63 as the outgroup. Both Lepidopteran and Dipteran species have evolved germline (purple box) and somatic (green box) paralogues. However, Lepidopteran germline gene underwent a further gene duplication to produce sperm-morph specific paralogues. For moth species, paralogues evolved apyrene-specific (red box) and eupyrene-specific (blue) functions. The monarch butterfly *D. plexippus* appears to have evolved two eupyrene-specific paralogues (blue text). Bootstrap values (100 repeats) are shown.

HCR-FISH revealed the expression patterns of *wompa* (LOC113515144; eupyrene) and *wimpa* (LOC113513840; apyrene) in *G. mellonella* larval and pupal testes. This confirmed a general pattern of high expression of *wompa* in larval spermatocytes, and high expression of *wimpa* in pupal spermatocytes, as expected from phylogenetic analysis (Fig. 6). The clear upregulation of *wimpa* in pupal testes vs larval testes also corroborated the RNA-seq data (Table S4, S5). HCR-FISH revealed *wimpa* and *wompa* co-expression in a small number of pupal cysts, predicted to be spermatocyte cysts committed to the eupyrene pathway (Fig. 6). *wompa* was relatively highly expressed across all primary spermatocyte samples in our RNA-seq dataset, with higher variation between pupal spermatocyte cyst samples (Table S5). This could indicate that both *wompa* and *wimpa* paralogues are important in later stages of eupyrene sperm development, whilst *wimpa* alone is necessary for apyrene sperm development.

**Figure 6.**
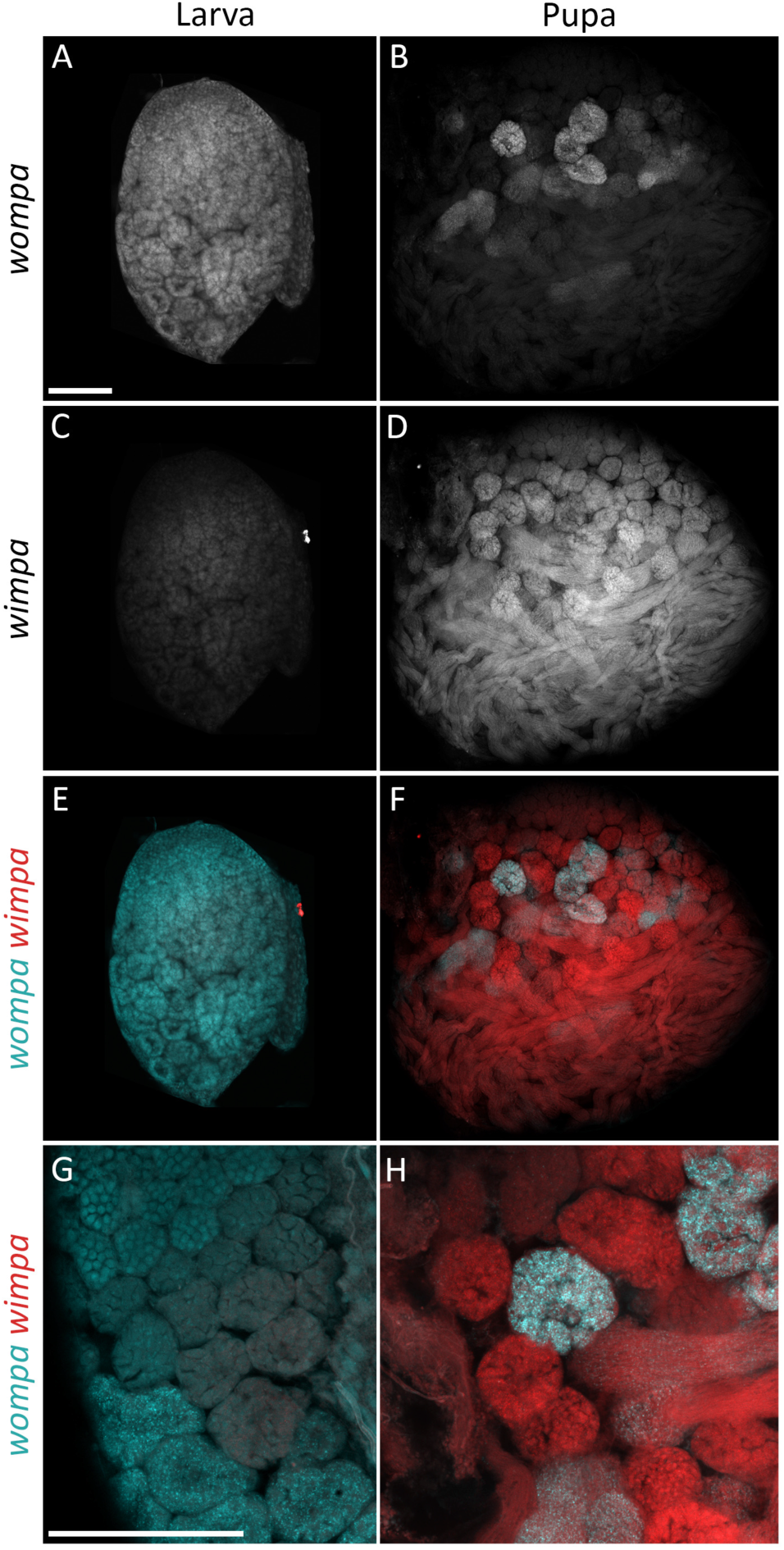
Differential expression of *Ccdc63* paralogues *wimpa* and *wompa* between developing apyrene and eupyrene sperm in *G. mellonella*. HCR-FISH analysis of *wompa* LOC113515144 (A, B; cyan in merged images E, F, G, H) and apyrene-enriched *wimpa* LOC113513840 (C, D; red in merged images E, F, G, H) of larval (A, C, E, G) and pupal (B, D, F, H) testes. High *wompa* and low *wimpa* expression was detected in larval eupyrene-destined spermatocytes (A, C). *wimpa* was highly expressed in spermatocytes and spermatids on the apyrene differentiation pathway in pupal testes (D). Predicted eupyrene-committed cysts in pupal testes demonstrated co-expression of both *wompa* and *wimpa* (large arrow), whilst predicted apyrene-committed cysts had *wimpa* expression only (arrowhead). Imaged using Zeiss LSM880 Airyscan upright confocal microscope. Scale bar is 100μm.

### Divergent expression of β*-tubulin* family members in Lepidoptera

From our RNA-seq data, another gene significantly upregulated in pupal spermatocytes was a β-tubulin gene (LOC113522729) of unknown genealogy (Table S4). Many β-tubulin genes (e.g. LOC113519435) were also amongst the most highly expressed in the spermatocyte transcriptomic data (Supplementary Data File 3). β-tubulin proteins, along with α-tubulin, constitute the microtubule cytoskeleton, which acts as a structural framework in cells; crucial for cell morphology, cell division, intracellular transport and the axoneme. In *D. melanogaster*, there are five β-tubulin genes. β*2* (β*-tubulin 85D)* is exclusively expressed in the male germline, is required for successful meiosis and axoneme elongation in spermatogenesis, and is abundant in the sperm proteome (28, 30, 31). β*-tubulin 65B* is also expressed exclusively in the male germline, but its role in spermatogenesis has not been determined, and the protein has not been detected in the sperm proteome (28). β1 (β*-tubulin 56D*) is expressed in both soma and male germline, and detected in the sperm proteome, while expression of both β3 (β*-tubulin 60D*) and β4 (β*-tubulin 97EF*) is restricted to the soma. *Aedes aegypti* has a similar β-tubulin gene tree, but has evolved two β4-tubulin paralogues (32). In *Bombyx mori*, four β-tubulin family members have described, including two somatic β1-tubulin paralogues (β1a and β1b), testis-specific β2-tubulin and somatic β3-tubulin. Whittington *et al.* (2019) (10) identified six β-tubulin proteins in the mature sperm proteomes of both *M. sexta* and *D. plexippus,* with differing sperm morph specificity. In *M. sexta,* one β-tubulin protein was apyrene-enriched while the remaining β-tubulin proteins were detected in both sperm morphs. Contrastingly, in *D. plexippus,* two eupyrene-specific β-tubulin proteins were identified, alongside one apyrene-specific β-tubulin protein and three shared proteins. We used phylogenetic analysis to resolve the evolutionary relationships of the sperm-morph specific β-tubulin proteins detected by Whittington *et al* (2019) (10), and the apyrene-specific β-tubulin *G. mellonella* gene.

A neighbour-joining phylogenetic tree of β-tubulin protein sequences is shown in Figure 7. Identification of the major subfamilies was via published assignments and confirmed by analysis of the C-terminal sequences (Fig. S3). Before the divergence of Lepidoptera and Diptera, a duplication of the ancestral β*4-tubulin* gene (green circle, Fig. 7), produced β*4-tubulin* (green box, Fig. 7), and β*4B-tubulin* (red box, Fig. 7). Subsequent Lepidopteran-specific gene duplications expanded the β4*-tubulin* family. The apyrene-specific β-*tubulin* gene (LOC113522729) from our RNA-seq (red asterisk, Fig. 7) is a semi-orthologue of the poorly characterised, germline-specific β*-tubulin 65B* from *D. melanogaster* (CG32396); both are β*4B-tubulins*. Whilst many β*4-tubulin* genes had low expression in the male germline, almost all the Lepidopteran β*4B-tubulin* paralogues were highly expressed in the male germline (10, 33) (red box, Fig. 7). The exception was *M. sexta* β*4B-tubulin* (LOC115455595), which was assumed to be somatic or lowly expressed due to absence from the sperm proteomic dataset. In contrast to *wimpa/wompa*, the β*4B-tubulin* paralogue lineages in Lepidoptera have not evolved clear apyrene– or eupyrene-enriched expression patterns, suggesting that the β*4B-tubulin* paralogues have independently acquired specific functions in sperm morph differentiation.

**Figure 7.**
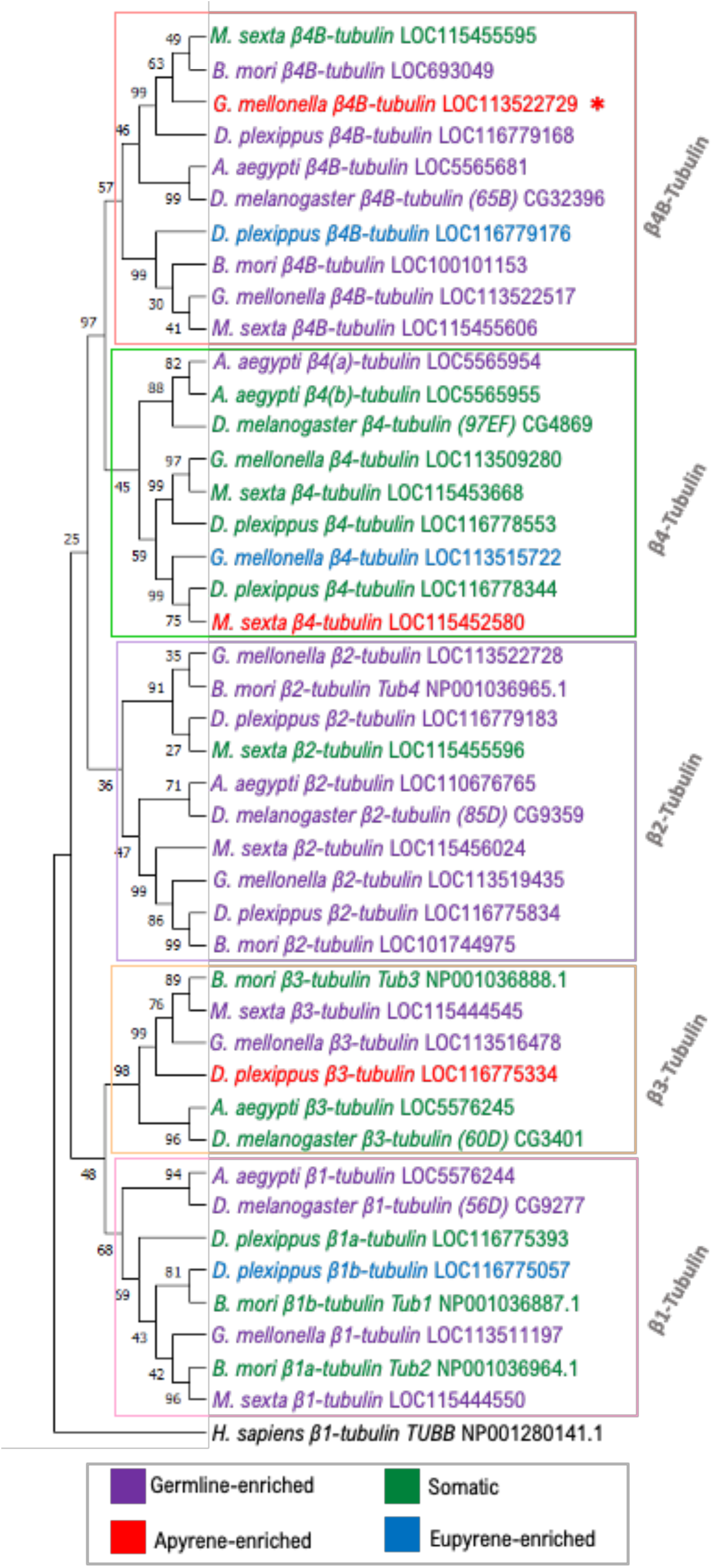
Phylogenetic analysis of the Lepidopteran β-tubulin family. Neighbour-joining tree of 42 β-tubulin proteins from Lepidoptera (*G. mellonella, D. plexippus, M. sexta, B. mori*) & Diptera (*D. melanogaster, A. aegypti*), with human (*H. sapiens*) β1-tubulin as the outgroup. Boxes outline the different β-tubulin family members. The β-tubulin gene significantly upregulated (p<0.05) in *G. mellonella* pupal spermatocytes is indicated by the red asterisk. Text colours denote expression pattern based on available transcriptomic and proteomic data: Apyrene-enriched (red), eupyrene-enriched (blue), germline-enriched (purple) or somatic (green). Bootstrap values (10,000 repeats) are shown.

*Β1-tubulin* and *Β3-tubulin* family members in Lepidoptera also demonstrated a divergence in expression patterns between Lepidopteran species (Fig. 7, Supplementary Data File 3). In contrast, almost all β*2-tubulin 85D* orthologues were expressed in the germline, with *G. mellonella* β*2-tubulin 85D* genes (LOC113519435, LOC113522728) showing very high expression levels in all spermatocyte samples (purple cluster, Fig. 7, Supplementary Data File 3). Phylogenetic analysis suggested that there are two β*2-tubulin 85D* paralogues in Lepidoptera, however the bootstrap value is relatively low for this gene duplication (purple circle, Fig. 7).

HCR-FISH analysis confirmed the results from the RNA-seq analysis. β*2-tubulin 85D* orthologue (LOC113519435) expression was detected at very high levels in spermatocytes and spermatids in both larval and pupal testes, suggesting that it is required for differentiation of both sperm morphs (Fig. 8A &B, Table S5). In contrast, *in situ* staining of apyrene-specific β4B-tubulin (LOC113522729) in *G. mellonella* testes revealed increased expression in pupal spermatocytes and spermatids vs larval spermatocytes (Fig. 8C & D, Table S5). A low expression level of the predicted apyrene-specific β4B-tubulin (LOC113522729) was detected by HCR-FISH in a subset of eupyrene primary spermatocyte cysts in the larval testes, with RNA-seq analysis also detecting a very low level (Fig. 8C, Table S5). This suggests that β*4B-tubulin* (LOC113522729) is of particular importance in apyrene sperm differentiation in *G. mellonella*. Overall, phylogenetic and expression analysis of β*-tubulin* genes revealed Lepidopteran-specific gene duplications which have not been previously identified. These β*-tubulin* duplications generated a suite of genes available for specialisation for different sperm morphs, but there was surprising variability of expression patterns of paralogues with respect to sperm morph between different moth and butterfly species.

**Figure 8.**
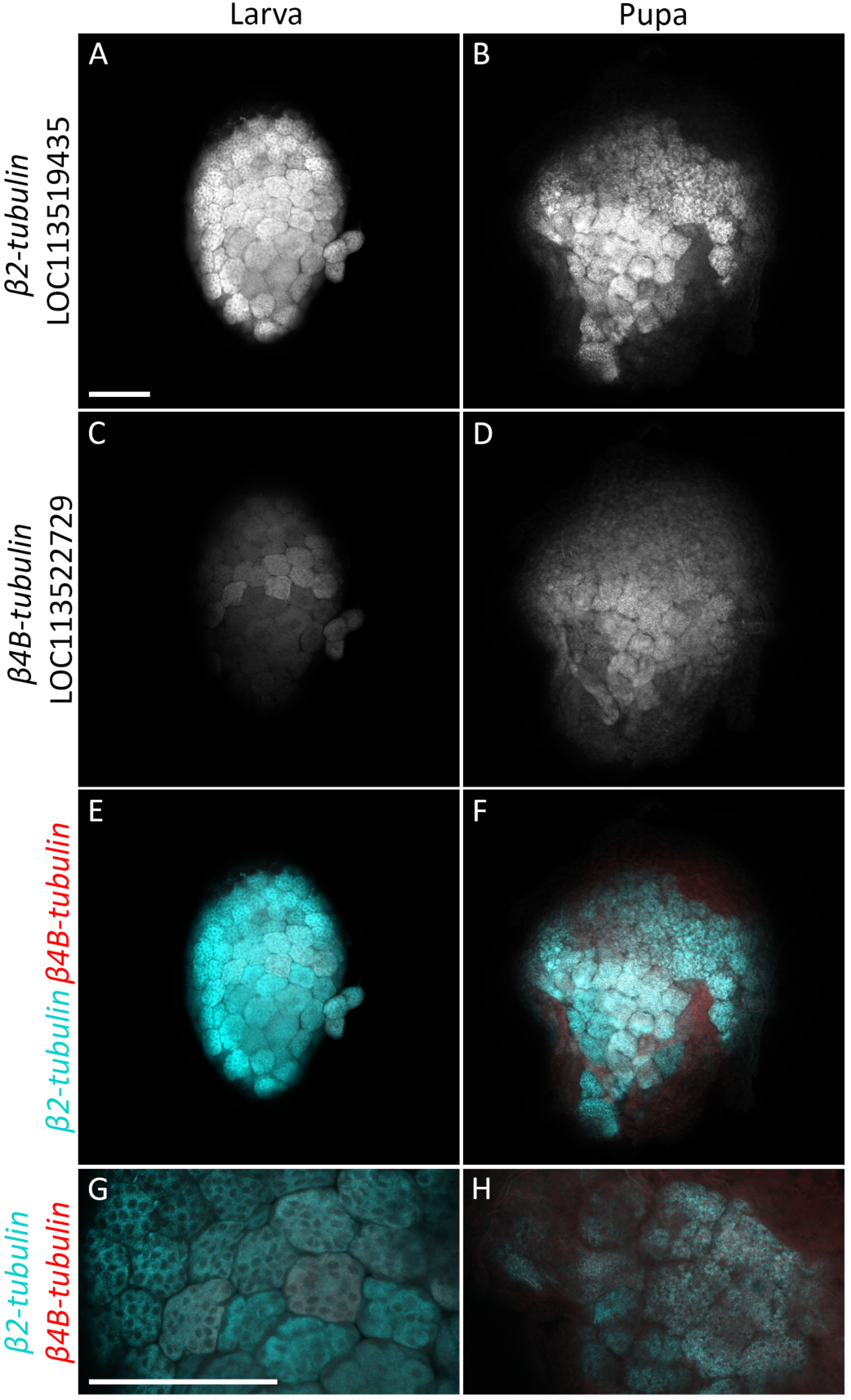
Ubiquitous vs apyrene-enriched expression of β*2-tubulin* and β*4B-tubulin* in *G. mellonella* testes. HCR-FISH analysis of β*2-tubulin* LOC113519435 (A, B; cyan in merged images, E, F, G, H) and apyrene-enriched β*4B-tubulin* LOC113522729 (C, D; red in merged images E, F, G, H), of larval (A, C, E, G) and pupal (B, D, F, H) testes. High levels of β*2-tubulin* were detected in both larval and pupal testes, corroborating ubiquitous germline expression (A, B). β*4B-tubulin* expression was higher in pupal testes vs larval (C,D), with low β*-tubulin* expression detected in a subset of eupyrene-destined primary spermatocytes in larval testes (C). Imaged using Zeiss LSM880 Airyscan upright confocal microscope. Scale bar is 100μm.

## Discussion

Sperm heteromorphism is present in almost all Lepidoptera, and the production of two sperm morphs is essential for fertility of moths and butterflies (1, 5, 6). The non-fertilising morphs are generated through precise, regulated processes, rather than through a variety of defective deviations from “normal”. Whilst the proteomes of the sperm morphs contain many shared proteins, eupyrene and apyrene sperm also contain unique, specialised proteins (10). How these differences are established earlier in spermatogenesis has not previously been described. In principle, production of two similar, but distinct, final cell morphologies can depend on genes i) expressed in both lineages, with the same timing but different absolute levels; ii) expressed exclusively in one or other lineage; iii) expressed in both lineages but with different temporal dynamics; iv) duplicated and subfunctionalised, such that the required protein function is provided by distinct isoforms. We validated examples of i (*Taf4*), iii (*sxl*) and iv (*wompa/wimpa*, and β*-tubulin*). With our sequencing data we cannot be sure of genes *exclusively* in one or other lineage, however we did find examples of single copy genes with dramatic differences in absolute expression level between cyst types. LOC113516308, a gene conserved across arthropods, with no known or predicted function had >16 fold higher expression in larval spermatocytes than pupal spermatocytes. Meanwhile, LOC116412852, which encodes a predicted plasma-membrane associated CAP-domain containing protein conserved across Lepidoptera, had >1000 fold higher expression in pupal spermatocytes than larval spermatocytes.

Our data confirms differential transcription of many genes, thus confirming that the unique sperm morphology is underpinned by differential transcription at the spermatocyte stage. Higher or lower expression of the general transcription factor Taf4, along with differential expression of other transcriptional regulators, may be implicated in establishing the distinct transcriptomes. Persistence of *Sxl* expression in the apyrene spermatocytes could affect RNA stability or could result in production of alternative splice variants in these cells. Among the DEGs we found several that could be regulate or enact the alternative meiosis seen in the apyrene differentiation programme (e.g. spindle proteins, cell cycle checkpoint proteins, meiotic recombination factors). Additionally, our DEG list includes many more examples of genes likely implicated in the differential elongation and final morphology. Validation of our RNA-seq dataset via phylogenetic analysis and identification of orthologous genes in published proteomic/ transcriptomic datasets demonstrates the predictive nature of early transcriptomic differences on the final proteomes and thus morphology of the mature sperm.

Phylogenetic analysis revealed the expansion of gene families in Lepidoptera via gene duplications, allowing for the paralogues to adopt specialised functions in producing the sperm heteromorphism phenotype. For example, duplication of the ancestral sperm axoneme component *Ccdc63* (27), produced two paralogous genes with distinct expression patterns in *G. mellonella* larval and pupal testes. They are predicted to play unique roles in eupyrene and apyrene sperm development, and have been termed *wompa* and *wimpa*, respectively. Ccdc63 has been duplicated to generate paralogues specifically for somatic axonemes and sperm axonemes in both Lepidoptera and Diptera (27), in addition to the sperm morph duplication in Lepidoptera. It may provide subtly different, but evolutionarily important, functionality to these distinct motile cilia, and comparative functional studies could be very interesting in the future.

A similar expansion of the well-known β*-tubulin* gene family was also observed in the Lepidopteran species studied. Lepidopteran-specific gene duplications of ancestral β*4B-tubulin* and β*2-tubulin* genes has led to many germline-enriched β*-tubulin* genes. We, again, predict that these paralogues have undergone sub-functionalisation to play key roles in sperm heteromorphism evolution, and in the normal function of the different morphs. Although there is evidence for sperm-morph enriched expression for a small number of β*-tubulin* genes, overall β*-tubulin* paralogues have not evolved a specific bias towards fertile or infertile sperm development, as found for *wimpa/wompa* paralogues. This suggests that exact function of the β*4B-tubulin* and β*2B-tubulin* paralogues in enacting or regulating dichotomous spermatogenesis varies between Lepidopteran species, revealing evolutionary flexibility in the co-option of genes in the process.

Interestingly, our RNA-seq analysis found a higher number of upregulated genes in pupal spermatocytes compared to larval, which contrasts to previous findings that apyrene sperm possess a *less* diverse proteome (10). While some of the differences could be species-specific, we note that Whittington et al. (2019) investigated mature sperm proteomes, whilst we have focused on the earlier spermatocyte transcriptome (10). During dynamic processes, such as differentiation, there may be only a moderate correlation between transcript and protein levels due to post-transcriptional regulation, providing a possible explanation for the observed disconnect (34). Increased expression of regulatory proteins, that are required at higher levels in the apyrene spermatocytes during spermatogenesis, but are not included in the final mature sperm, could also contribute to the observed pattern. Furthermore, we hypothesise that the higher transcriptomic diversity of developing apyrene sperm may make evolutionary sense in an extension of the “out of the testis” phenomenon (35); apyrene spermatocytes may provide a playground to experiment with newly evolved genes in an environment with lower functional constraints in comparison to the fertilising eupyrene sperm. Differential signatures of selection have already been described for proteins differentially expressed between morphs, with apyrene sperm-specific proteins showing little evidence of positive selection (36).

Our Lepidopteran model of sperm heteromorphism, the wax moth *Galleria mellonella*, is an emerging model organism within the life sciences, predominantly as a model species for investigating the response to infection (37). Genetic tools such as transgenesis and CRISPR-Cas9 genome editing are actively being developed in *Galleria mellonella*, making it an attractive model species for future research into sperm heteromorphism regulators (38).

Fundamental research into Lepidopteran reproduction has potentially wide-reaching implications in terms of controlling Lepidopteran pests in agriculture. Lepidopteran pests cause significant economic damage due to crop loss, with the problem increasing due to globalisation causing further spread of invasive Lepidopteran species (39). The wax moth *Galleria mellonella* is a significant pest of honeybees (40). With the importance of biodiversity becoming increasingly prevalent in the public consciousness, innovative solutions are required to effectively target these pest populations. Seth *et al.* (2023) recently proposed the production of infertile apyrene sperm as a potential ‘Achilles heel’ to specifically target Lepidopteran pests (2). Our data set provides a resource to mine to investigate novel genes implicated in sperm differential morphogenesis, and specifically target genes involved in apyrene sperm development. On the other hand, improved understanding of Lepidopteran reproduction could be important in the broader context of declines in Lepidopteran pollinator populations (41). The possible future decline in fertility and hence survival of moth and butterfly pollinator populations due to these environmental factors may have devastating impacts on our ecosystems and global food production.

## Data availability

The RNAseq data sets are available on NCBI SRA, accession number PRJNA1028403.

## Acknowledgments

We thank the Galleria Mellonella Research Centre, University of Exeter (www.gmrcuk.org), especially James Wakefield and James Pearce, for *G. mellonella* larvae and food, and for initial training in dissection, and helpful discussions through the project. We thank Henry Krause and Catherine Shao for training in HCR-FISH, and Matthew Jachimowicz for the HCR probe design script. We thank the Cardiff University *Drosophila* research labs for discussions, and Pete Kille for critical reading of the manuscript. Sample preparation, quality control and RNA sequencing and data processing was performed with support from the Cardiff University School of Biosciences Genome and Biocomputing hubs, and we are especially grateful to Angela Marchbank for her input; imaging was supported by the Biosciences Imaging hub. This work was supported by Cardiff University and the Biotechnology and Biological Sciences Research Council [grant number BB/T006129/1].

## Supplementary Methods

### 1. G. mellonella culture

Wild type *G. mellonella* larvae were stored in the dark at room temperature and maintained on food medium based on diet 3 of (1), consisting of 250 g cornmeal, 150 g yeast extract, 100 g soy flour, 100 g powder milk, 200 g honey, 200 g glycerol and beeswax blocks.

### 2. Fluorescence Hybridisation Chain Reaction (HCR-FISH)

HCR-FISH v3 was used to visualise gene expression (2). Probe pairs against target *G. mellonella* mRNA sequences obtained from NCBI were designed using a python script with a specified hairpin amplifier sequence. The specificity of the probe pairs was assessed using NCBI nucleotide blast. Final *G. mellonella* probe sequences are in Supplementary Table 2.

*G. mellonella* larvae and pupae were collected into a petri dish on ice, and larvae only were additionally anaesthetised using CO_2_. *G. mellonella* testes (n=10) per sample were dissected in PBS, ensuring removal of all fat body, and incubated in 1 ml 0.5% paraformaldehyde (PFA) in PBT buffer (1x PBS, 0.1% Tween 20) at room temperature (RT) for 10 minutes. PFA 0.5% was removed and replaced with 4% PFA in PBT buffer and incubated on the nutator for 30 minutes. Samples were washed twice with PBT for 5 minutes on the nutator. Samples were rinsed with methanol, and then stored in fresh methanol at –18°C. Testes were washed with PBT after storage in methanol, and incubated in 100 µl 30% Hybridisation buffer (Hyb, 30% formamide, 5X SSC, 9 mM citric acid (pH 6.0), 0.1% Tween 20, 50 μg/mL heparin, 5X Denhardt’s solution, 10% dextran sulphate) at 37°C for 30 minutes. Per gene, 3.2 µl of 100µM DNA oligonucleotide probes (Integrated DNA Technologies) were added to 100 µl Hyb and incubated at 80°C for 5 mins. Testes were simultaneously incubated in Hyb at 80°C for 5 mins. The Hyb for each sample was replaced with the appropriate Hyb & probe solution and incubated for a further 5 mins at 80°C, and then incubated at 37°C overnight. The Hyb & probe solution was replaced and testes washed with 30% probe wash buffer (30% formamide, 5X SSC, 9 mM citric acid (pH 6.0), 0.1% Tween 20, 50 μg/mL heparin) at 37°C. Fluorescently labelled DNA (2µl of each) hairpin amplifier (3 pmol/μl, H1, H2) with B1, B2, B3 or B4 sequences (Molecular instruments) were heated to 95°C for 90 seconds and then cooled to RT. Hairpin pairs were added to amplification buffer (5X SSC, 0.1% Tween 20, 10% dextran sulphate) in a final volume of 50 µl, and the testes were incubated with gentle rocking at RT overnight. Amplifier mix was removed, and samples washed with 4x 5 minutes with SSCT wash buffer (5X SSC, 0.1% Tween 20), and then subsequently washed in 3x 5 minutes in PBS. A subset of samples were incubated in 1 µg/ml Hoescht (33258) in PBS to stain DNA. Samples were mounted for lightsheet microscopy analysis in 1% low melting point agarose in PBS (Sigma) within 1.5 mm glass capillary tubes using plungers.

Fluorescence was visualised using Zeiss Lightsheet Z.1 system in the Cardiff Bioimaging Hub. A 20X detection objective, and two 10X illumination objectives were used, in conjunction with a water chamber filled with PBS. Acquisition settings were: 16-bit with dual-side fusion and pivot scan. 561 nm laser was used to visualise Alexa546 amplifiers, and 488n m laser to visualise Alexa488 amplifiers. Z-stacks were obtained of the whole testis sample and subsequently analysed in Arivis vision 4D 2.12.3 x64 software to produce maximum brightness projections. Alternatively, samples were mounted on slides in 85% glycerol, 2.5% n-propyl gallate, and imaged using a Zeiss LSM880 Airyscan upright confocal microscope with X40 oil objective.

### 3. Dissection of *G. mellonella* testes and isolation of spermatocyte cysts for RNA-seq

Five 6^th^ instar larvae and five three-day old pupae were collected into a petri dish on ice, and larvae were additionally anaesthetised using CO_2_. Testes were dissected in testis buffer (183 mM KCl, 47mM NaCL, 10 mM Tris HCl, pH6.8) to obtain a single follicle. Follicle contents were spilled onto a siliconised slide, and contents visualised using Nikon Eclipse Ti-S inverted microscope. Two primary spermatocyte cysts per follicle were selected based on size/morphology and imaged at 40X magnification using Hamamatsu ORCA-05G camera attachment and HCImage software (v.2,2,6,4 Hamamatsu Corp. 2011). Selected cysts were collected into separate 0.5 ml tubes using stretched glass Pasteur pipettes and immediately frozen using liquid nitrogen and stored at –80°C until use.

### RNA-seq library preparation

RNA libraries were produced from the 20 primary spermatocyte cysts using the QIAseq FX Single Cell RNA Library Kit (Qiagen). Following cell lysis and gDNA degradation, mRNA was extracted and amplified from the samples using Qiagen’s protocol ‘Amplification of Poly A+ mRNA from Single Cells’. cDNA product of this step was stored at –20°C. The cDNA was then converted into libraries following the Qiagen protocol ‘Enzymatic Fragmentation and Library Preparation Using QIAseq FX Single Cell Amplified cDNA’. cDNA concentration was quantified using a Qubit assay (ThermoFisher Scientific), following the manufacturer’s protocol. Library quality and fragment size was assessed using capillary electrophoresis in the D1000 Tapestation system (Agilent). cDNA library was pooled to an equimolar concentration of 3 nM according to the molarity of the desired 150 bp to 600 bp fragment sizes from the Tapestation analysis. The pooled cDNA was subsequently cleaned and concentrated to 13 ng/µl in a final volume of 35 µl by the Cardiff Genomics Research Hub team, and DNA size selection was carried out using the Blue Pippin system on a 2% agarose cassette (Sage Science). Sequencing was completed using the Illumina NextSeq500 Sequencer at Cardiff University Heath hospital site, and de-multiplexed by the Cardiff Genomics Research Hub.

### 5. RNA-seq bioinformatics and statistical analysis

Sequence data quality was assessed using FastQC (3) and poor-quality sequences and adapters were trimmed using Trimmomatic tool v0.39 (4). Sequence alignment was performed using Hisat2/2.2.1 (5), with the transcript sequences aligned to the *G. mellonella* reference genome (CSIRO_AGI_GalMel_v1, Rahul Vivek Rane 2022) from NCBI (6). SAM files were converted into sorted BAM files with Samtools v1.15 (7). FeatureCounts from the subread v2.0.2 package (20) was used to count how many reads mapped to genomic features (8).

### 6. Phylogenetic analysis of DEGs of interest

BLASTP output of *G. mellonella* homologues from species of interest were aligned using ClustalW with default settings in MEGA V. 11.0. 13 software (9). Redundant protein isoforms, partial sequences and hypothetical proteins were removed from analyses. For Ccdc63, the alignment output was input into Gblocks with low stringency settings (10). A maximum-likelihood tree using the LG substitution model was produced with 100 bootstrap repeats in MEGA. For β-tubulin analysis, Gblocks analysis was not required due to the highly conserved nature of β-tubulin proteins. A neighbour-joining (Poisson model) phylogenetic tree was produced with 10,000 bootstrap repeats using MEGA.

## Supplementary data

**Supplementary Table 1:** Summary of previously discovered genes with roles in sperm heteromorphism regulation in Lepidoptera, and corresponding RNA-seq results from *G. mellonella* cysts. Previously described spermatogenesis or sperm heteromorphism regulators in other Lepidopteran models (*Bombyx mori*, *Spodoptera litura*). For each gene, RNA-seq results from our *G. mellonella* dataset comparing larval and pupal primary spermatocyte transcriptomes are shown, including Log_2_ Fold change (Log_2_FC), adjusted p-value, and average read count across all samples (larval and pupal).

| Gene | Protein function | Spermatogenesis role determined in other lepidoptera | Ref(s) | ***G. mellonella* Primary Spermatocyte RNA-seq results** |  |  |  |
| --- | --- | --- | --- | --- | --- | --- | --- |
|  |  |  |  | Gene ID | Log<sub>2</sub>FC (pupal vs larval) | Adj. P-value | Average read count |
| *BmSex-lethal (BmSxl)* | RNA-binding protein. | *B. mori* - apyrene<br>*S. litura* –eupyrene and apyrene | (11)<br>(12) | LOC113515001 | 1.475 | 0.08 | 3544 |
| *BmAly (Lin9)* | Transcription complex component | *B. mori* –eupyrene. Role in apyrene not investigated. | (13) | LOC113523285 | -0.07 | 0.95 | 690 |
| *BmPNLDC1* | Pre-piRNA trimmer. | *B. mori* – Eupyrene | (14,<br>15) | LOC113517413 | -0.196 | 0.84 | 14 |
| *BmHen1* | Small RNA 2'-O-methyltransferase of piRNAs. | *B. mori* – Eupyrene | (15) | LOC113522915 | -0.732 | 0.24 | 2465 |
| *BmPMFBP1* | Polyamine modulated factor 1 binding protein 1. | *B. mori* – Eupyrene | (16) | No orthologue | - | - | - |
| *BmMaelstrom* | RNA binding protein. | *B. mori* – Apyrene and Eupyrene | (17) | LOC113521157 | -0.631 | 0.416093 | 12 |
| *BmPrmt5* | Type II arginine methyltransferase | *B. mori* – Apyrene and Eupyrene | (18) | LOC113522473 | 0.276 | 0.72 | 581 |
| *BmVasa* | ATP-dependent RNA helicase. | *B. mori* – Apyrene and Eupyrene | (18) | LOC113521823 | -0.441 | 0.59 | 570 |
| *BmSIWI* | piRNA binding protein | *B. mori* – Not required for spermatogenesis | (15,<br>19) | LOC113522979 | 0.85 | 0.30 | 520 |

**Supplementary Table 2:**
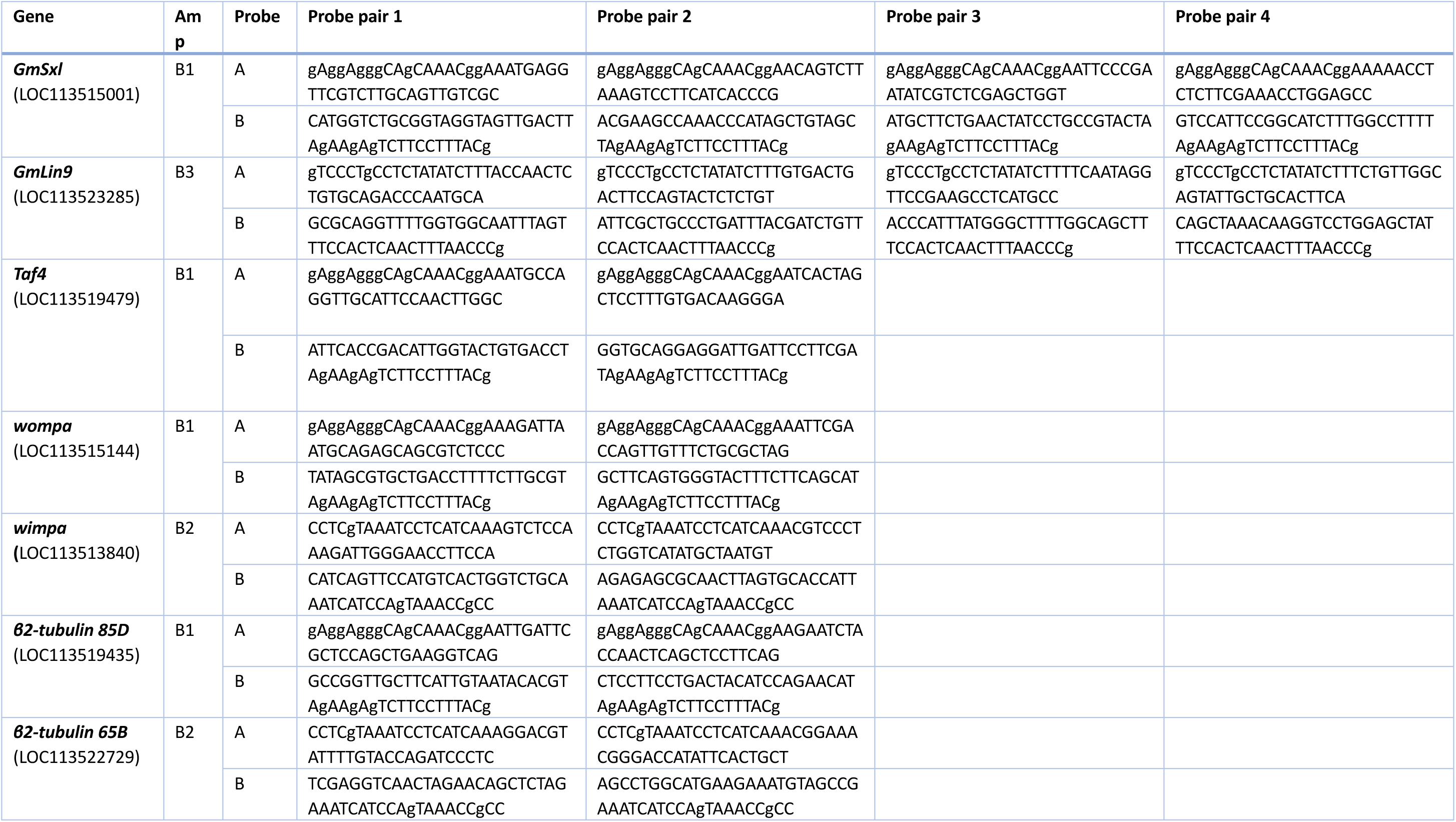
HCR-FISH probe pairs against target genes.

**Supplementary Table 3:** Sequencing depth, trimming and mapping of reads from RNA-seq of *G. mellonella* primary spermatocytes.

| Sample | Total raw paired end reads | Reads filtered by Trimmomatic (%) | Hisat2 input reads | Hisat2 uniquely mapped reads (%) |
| --- | --- | --- | --- | --- |
| Larv_1.1 | 9359490 | 7.19 | 8686427 | 79.95 |
| Larv_1.2 | 7720243 | 8.33 | 7077349 | 76.31 |
| Larv_2.1 | 11879451 | 10.29 | 10656613 | 61.59 |
| Larv_2.2 | 17328718 | 8.28 | 15894591 | 74.73 |
| Larv_3.1 | 19542755 | 9.46 | 17693934 | 73.09 |
| Larv_3.2 | 10313332 | 8.79 | 9406745 | 72.40 |
| Larv_4.1 | 12129493 | 9.80 | 10940328 | 71.91 |
| Larv_4.2 | 10087767 | 9.24 | 9155730 | 72.93 |
| Larv_5.1 | 16207682 | 11.27 | 14381523 | 60.28 |
| Larv_5.2 | 13671017 | 14.17 | 11733705 | 1.32 |
| Pupa_1.1 | 23316658 | 8.69 | 21291027 | 64.83 |
| Pupa_1.2 | 15942060 | 7.54 | 14740183 | 75.64 |
| Pupa_2.1 | 17180895 | 8.50 | 15720577 | 72.73 |
| Pupa_2.2 | 17970280 | 10.99 | 15995021 | 48.17 |
| Pupa_3.1 | 14626438 | 9.59 | 13224026 | 64.64 |
| Pupa_3.2 | 32138830 | 12.33 | 28177622 | 57.67 |
| Pupa_4.1 | 21544565 | 11.91 | 18978788 | 71.01 |
| Pupa_5.1 | 6456211 | 10.99 | 5746352 | 68.66 |
| Pupa_5.2 | 15062814 | 15.58 | 12715403 | 60.57 |

**Supplementary Table 4:** Summary of literature search and DAVID GO analysis to identify DEGs of interest from RNA-seq dataset. From the lists of significantly differentially expressed genes between larval and pupal primary spermatocytes, a subset of genes were elucidated as having possible roles in *G. mellonella* sperm heteromorphism regulation. Results of the literature search to investigate any previously found roles of DEGs in spermatogenesis, in Lepidoptera and other model organisms (i.e. mouse, *Drosophila melanogaster*). DAVID bioinformatic analysis identified enriched Gene Ontology (GO), UniprotKB Keywords (UP-KW) and Interpro terms in submitted gene lists. For each DEG, Log2 Fold change and adjusted P value are shown.

| Gene name & LOC | Short Name | Gene function | Associated enriched GO/UP-KW/INTERPRO term | Highest expression | Log2 FC | Adj. P-value | Reference |
| --- | --- | --- | --- | --- | --- | --- | --- |
| Transcription Initiation Factor TFIID subunit 4 (LOC113519479) | *Taf4* | TFIID subunit, transcription, spermatogenesis. Testis-specific paralogue *nht* required for testis transcriptional programme in *D. melanogaster* spermatocytes. | UP-KW-0804~Transcription | Larva | 2.57 | 0.0031 | (20) |
| Chromatin-remodelling ATPase INO80 (LOC113520261) | *Ino80* | Chromatin remodeller. Deletion in mice caused meiotic arrest. | GO:0016887~ATPase activity, UP-KW-0238~DNA-binding | Larva | 2.22 | 0.0312 | (21) |
| DNA mismatch repair protein Mlh1 (LOC113514204) | *Mlh1* | Promotes homologous recombination. Mutant male and female mice have unpaired univalent chromosomes leading to meiotic arrest and infertility. B. mori – co-promoting protein. | GO:0016887~ATPase activity | Larva | 2.68 | 6.27E-05 | (22) |
| DNA replication licensing factor Mcm6 (LOC113514226) | *Mcm6* | Initiates meiotic recombination. Deletion in mice caused meiotic arrest. | GO:0016887~ATPase activity, UP-KW-0238~DNA-binding | Larva | 2.23 | 0.0033 | (23) |
| Meiotic recombination protein SPO11 (LOC113517093) | *Spo11* | Initiates meiotic recombination. Deletion in mice caused meiotic arrest. Found in *B. mori* to be important for DSB formation. | - | Larva | 2.57 | 0.0176 | (22, 24) |
| E3 ubiquitin-protein ligase UHRF1-like (LOC113516552) | *Uhrf1* | Multidomain protein; regulator of many epigenetic pathways. Conditional deletion in mice leads to meiotic arrest and male infertility. | UP-KW-0238~DNA-binding | Larva | 1.76 | 0.0019 | (25) |
| Inhibitor of growth protein 5(LOC113516070) | *Ing5* | Putative tumour suppressor. 5 family members (1-5). Spermatocytes in mice deficient in family member *Ing2* underwent meiotic arrest, leading to infertility. | GO:0008285~negative regulation of cell proliferation | Larva | 2.20 | 0.007957 | (26) |
| Coiled-coil domain containing protein 63 (LOC113513840) | *Ccdc63* | Component of outer dynein arm docking complex, involved in formation of the sperm axoneme. Required for male fertility in *D. melanogaster*. | UP-KW-0175~Coiled coil | Pupa | 7.16 | 1.66E-14 | (27) |
| Tubulin beta chain-like (LOC113522729) | *Tbb* | Microtubule cytoskeleton; forms meiotic spindle. Flagellum formation. | - | Pupa | 9.09 | 8.37E-22 | (28) |
| Sperm flagellar protein 1-like (LOC113514483) | *Spef1* | Axoneme growth in spermiogenesis. Motile flagella. | GO:0031514~motile cilium | Pupa | 3.06 | 0.0003 | (29) |
| Cell division cycle protein 27 homolog (LOC113521294) | *Cdc27* | E3 ubiquitin ligase. Subunit of anaphase-promoting complex (APC/C) involved in regulating cell cycle progression. | UP-KW-0132~Cell division | Pupa | 2.15 | 0.0037 | (30) |
| Katanin p60 ATPase-containing subunit A1-like (LOC113515509) | *Katnal1* | Microtubule-severing enzyme. Katnal2 implicated in remodelling of mitotic/meiotic bipolar spindle. | UP-KW-0132~Cell division | Pupa | 2.134 | 0.0214 | (31) |
| Angiotensin-converting enzyme (LOC113518907) | *Ace* | Peptidyl dipeptidase. Implicated in apyrene sperm motility, *as in vitro* inhibition led to reduced *B. mori* apyrene sperm motility | UP-KW-0121~Carboxypeptidase | Pupa | 3.43 | 0.0001 | (32, 33) |
| Uncharacterised<br>LOC116412852 | - | Plasma membrane associated CAP-domain containing protein. CAP protein superfamily members have been implicated in male fertility in mammals. | GO:0016021~integral component of membrane;<br>IPR014044:CAP domain | Pupa | 9.326 | 5.76E-11 | (34) |

**Supplementary Table 5:** RNA-seq normalised counts for *G. mellonella* spermatogenesis genes validated by HCR-FISH and phylogenetic analysis.

| Gene | Sperm-morph enrichment | Log2FC | P-adj. | Larv_1<br>.2 | Larv_2<br>.2 | Larv_3<br>.1 | Larv_3<br>.2 | Larv_4<br>.1 | Larv_4<br>.2 | Pupa_1<br>.1 | Pupa_1<br>.2 | Pupa_3<br>.1 | Pupa_3<br>.2 | Pupa_4<br>.1 | Pupa_5<br>.1 | Pupa_5<br>.2 |
| --- | --- | --- | --- | --- | --- | --- | --- | --- | --- | --- | --- | --- | --- | --- | --- | --- |
| *GmSxl*<br>(LOC113515001) | Apyrene | 1.475 | 0.081399 | 376 | 1055 | 1810 | 681 | 1213 | 1232 | 21708 | 13240 | 655 | 1234 | 827 | 1987 | 62 |
| *GmLin9*<br>(LOC113523285) | Shared | 0.07 | 0.945633 | 272 | 2186 | 106 | 791 | 675 | 313 | 563 | 449 | 55 | 544 | 804 | 597 | 1614 |
| *Taf4*<br>(LOC113519479) | Eupyrene | 2.57 | 0.0031 | 606 | 280 | 132 | 338 | 380 | 923 | 60 | 191 | 25 | 11 | 12 | 146 | 20 |
| *wompa*<br>(LOC113515144) | Eupyrene | 0.027 | 0.97923 | 3070 | 3779 | 8642 | 991 | 5890 | 6877 | 44 | 696 | 8990 | 19679 | 3917 | 1091 | 1138 |
| *wimpa*<br>(LOC113513840) | Apyrene | 7.161 | 1.66E-14 | 66 | 6 | 4 | 47 | 48 | 39 | 24234 | 8214 | 5367 | 6757 | 2319 | 2106 | 93 |
| *β4B-tubulin*<br>(LOC113522729) | Apyrene | 10.098 | 1.71E-32 | 31 | 19 | 12 | 4 | 14 | 10 | 75555 | 26965 | 3474 | 15400 | 8349 | 6161 | 1584 |
| *β2-tubulin 85D*<br>(LOC113519435) | Shared | 0.217 | 0.802273 | 64596 | 57904 | 61697 | 58048 | 75195 | 64051 | 323154 | 76029 | 79670 | 31432 | 23672 | 38897 | 9610 |

**Supplementary Figure 1.**
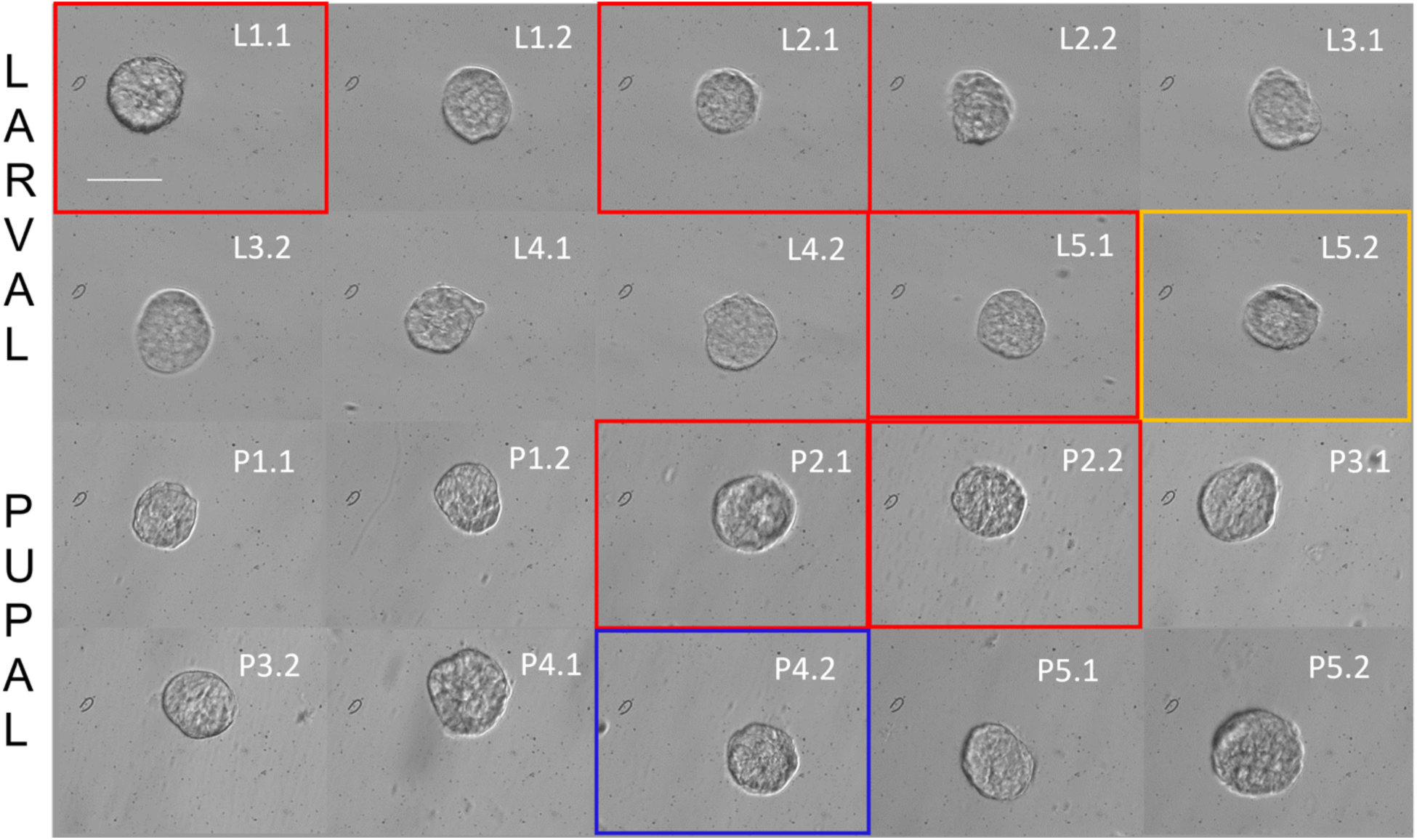
Primary spermatocyte cysts dissected from five *G. mellonella* larvae (L1-L5) and five pupae (P1-P5) for RNA-seq. cDNA libraries were subsequently prepared from cysts for RNA-seq. Red boxes show primary spermatocyte cysts excluded from DEG analysis due to ambiguous clustering. Larval cyst 5.2 was excluded from analysis due to very low hisat2 mapping % (orange box). Pupal cyst 4.2 was not sequenced due to low cDNA concentration (blue box). Scale bar = 60µm.

**Supplementary Figure 2.**
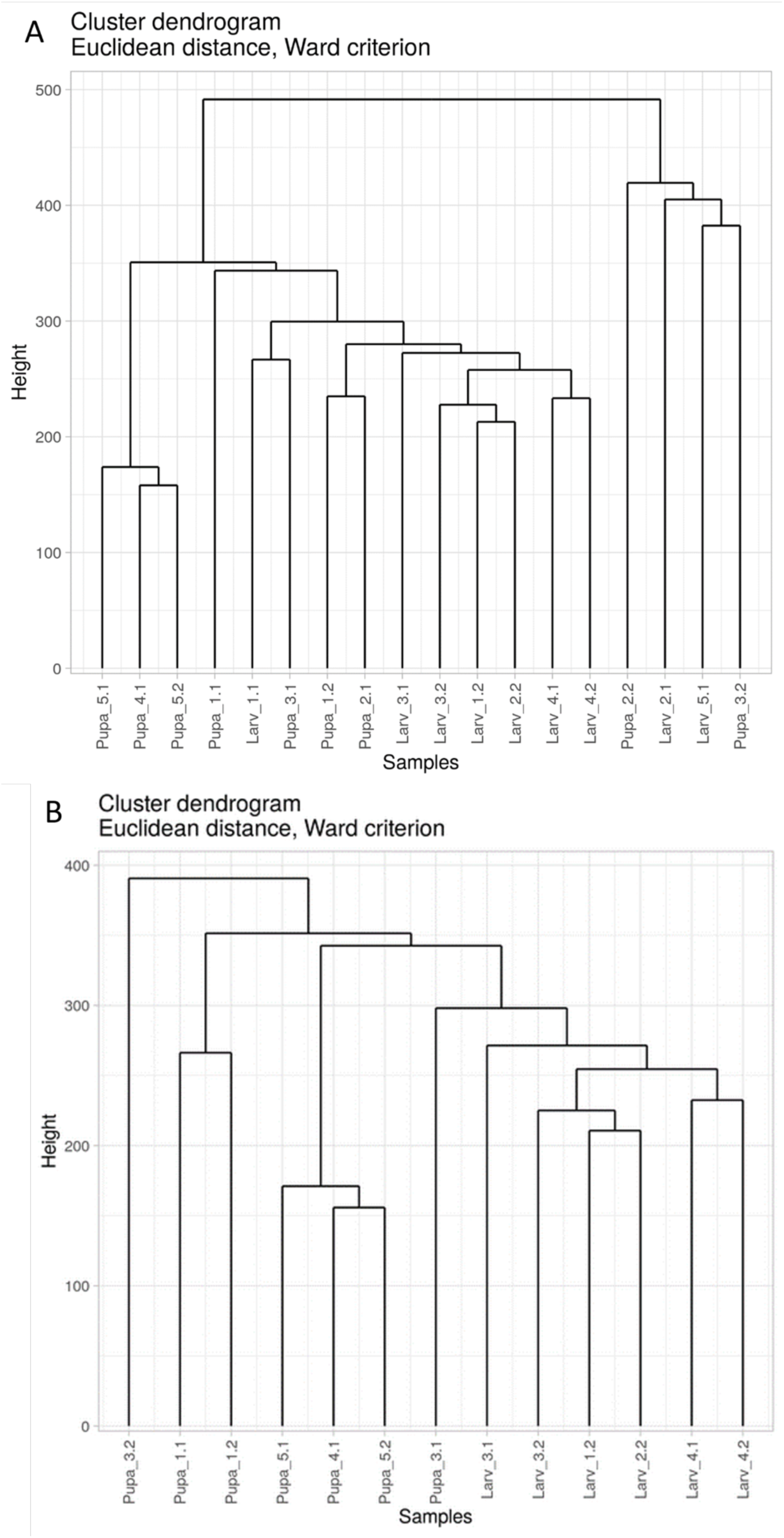
Hierarchal dendrogram of all samples and after filtering for larval and pupal clusters. Hierarchal dendrograms were produced using SARTools R package (35) for A) all samples and B) remaining samples after filtering out ambiguous samples.

**Supplementary Figure 3:**
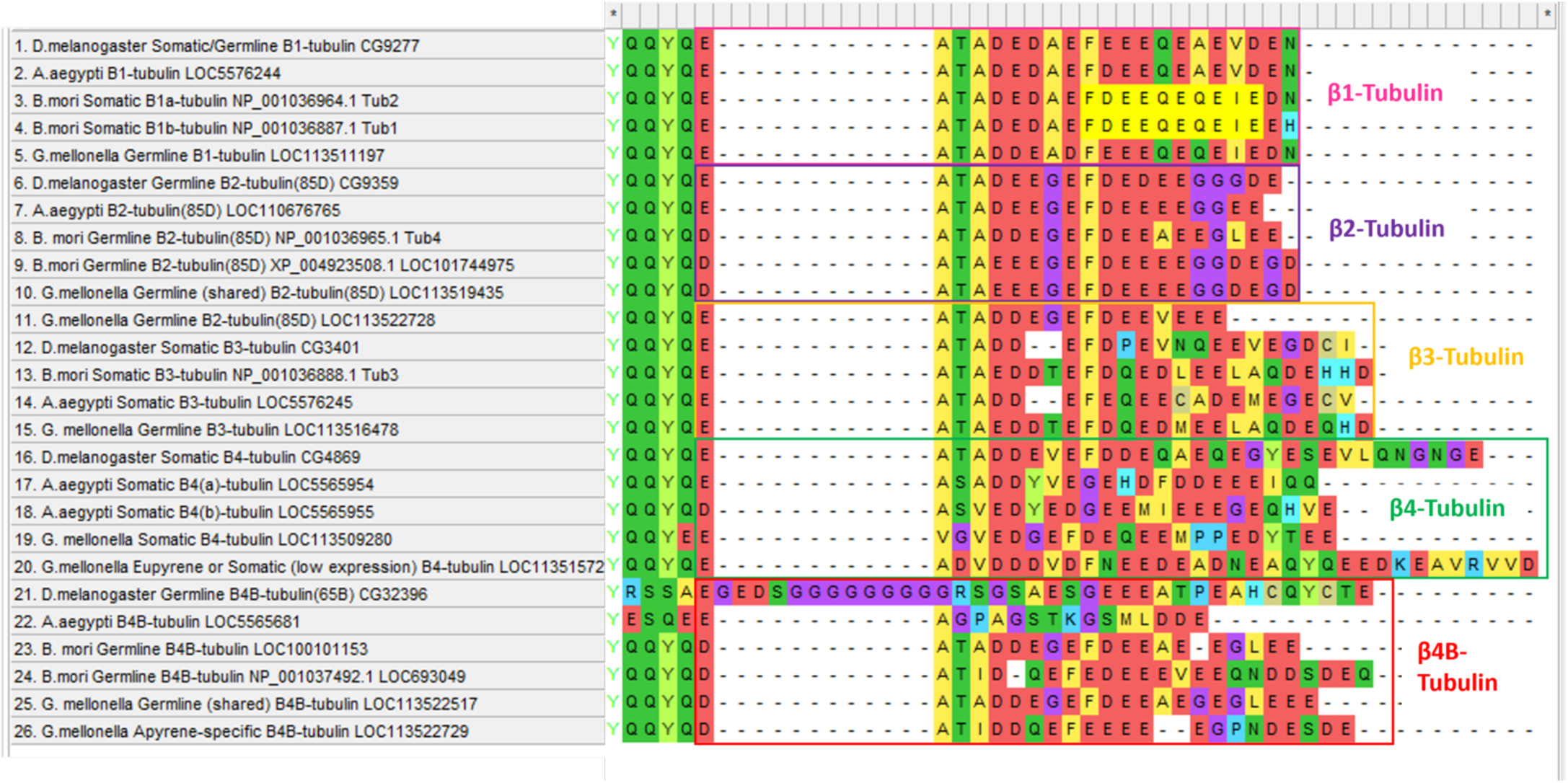
β-tubulin C-terminus protein sequence alignment to identify β-tubulin family members. MEGA ClustalW alignment of *D. melanogaster, A. aegypti, B. mori* & *G. mellonella* β-tubulin proteins, with only the variable COOH terminus ends shown (amino acid 438 onwards). The C-terminus sequence varies between β-tubulin family members (β1, 2, 3, 4 & 4B), but is relatively highly conserved within the same orthologous branch. β-tubulin CTD sequences of *B. mori, D. melanogaster* and *A. aegypti* from Table 1 of (28) were used to validate our β-tubulin orthologues.

